# Soft ticks perform evaporative cooling during blood-feeding

**DOI:** 10.1101/2020.06.30.180968

**Authors:** Claudio R. Lazzari, Aurélie Fauquet, Chloé Lahondère, Ricardo N. Araújo, Marcos H. Pereira

## Abstract

Feeding on the blood of warm-blooded vertebrates is associated to thermal stress in haematophagous arthropods. It has been demonstrated that blood-sucking insects protect their physiological integrity either by synthesising heat-shock proteins or by means of thermoregulatory mechanisms. In this work, we describe the first thermoregulatory mechanism in a tick species, *Ornithodoros rostratus*. By performing real-time infrared thermography during feeding on mice we found that this acarian eliminates big amounts of fluid (urine) through their coxal glands; this fluid quickly spreads over the cuticular surface and its evaporation cools-down the body of the tick. The spread of the fluid is possible thanks to capillary diffusion through the sculptured exoskeleton of *Ornithodoros*. We discuss our findings in the frame of the adaptive strategies to cope with the thermal stress experienced by blood-sucking arthropods at each feeding event warm-blooded hosts.

## 1. Introduction

Many arthropod species have adopted the blood of endothermic vertebrates as a main or sole source of food. Blood is rich in nutrients and, except for the potential presence of pathogens, otherwise sterile. Yet, obtaining a blood meal and handling it is far from being an easy task for these animals. First, the blood flows inside vessels hidden under the skin of active and relatively much bigger animals, which are capable of defending themselves or even predate the aggressor, constitutes the first risky step of blood-feeding. Second, ingesting relatively huge amounts of hot blood, represents a major challenge for blood-sucking arthropods as it can lead to heat stress and challenges physiological balance (Benoit et al. 2019; Lehane 2005; Sterkel et al. 2017).

Among blood-sucking Acari, two different feeding strategies are found. While Ixodidae hard ticks attach to the host skin and slowly consume the host blood for long periods that last several days, Argasidae soft ticks ingest large amounts of blood in a relatively short time (McCoy et al. 2010). In contrast to Ixodidae, in which salivary glands play a major role in eliminating water ingested during blood meal (Kim et al. 2017), argasid ticks eliminate the excess of water in their blood meal *via* their coxal gland (McCoy et al. 2010; Sonenshine 2013).

In the present study, we focused on the soft tick *Ornithodoros rostratus* (Aragão, 1911) as experimental model. *O. rostratus* is an argasid tick that uses an eclectic array of food sources, including dogs, pigs, cows, peccaries and humans (Cançado et al., 2008; Hoogstraal, 1985; Ribeiro et al., 2013). These soft ticks are of medical and veterinary importance as they give a painful and itchy bite (Aragão, 1936; Estrada-Pena and Jongejan, 1999), due to their salivary compounds and telmophagous feeding mode, *i.e.* gathering the blood pooled in a provoked wound (Mans and Neitz, 2004). They may carry and transmit pathogens such as *Rickettsia rickettsii* and *Coxiella burnetii*, the causative agents of the Rocky Mountain spotted fever and acute Q fever, respectively, to humans and other animals (Almeida et al., 2012; Hoogstraal, 1985). This species of ticks has been reported in four countries in South America: Argentina, Bolivia, Brazil and Paraguay (Barros-Battesti et al., 2006; Guglielmone et al., 2003). During its life cycle, this species has one larval and three to six nymphal instars, and adults can arise from 3^rd^ to 6^th^-instar nymphs (Costa et al. 2015). This species is an obligatory hematophagous arthropod in all stages and remains in contact with its host for a period varying from a few minutes to hours (nymphal instars and adults), or even days (larvae) (Costa et al., 2015, 2016; Lavoipierre and Riek, 1955; Ribeiro et al., 2013). Although contact with their host is for a short period of time as compared to hard ticks, it remains crucial as their success in obtaining blood will then affect their survival, development and reproduction (Anderson and Magnarelli, 2008). Additionally, the shorter the contact, the lower is the risk of being perceived and killed by the host (Rossignol et al., 1985). Importantly, it is during blood feeding that pathogens circulate between vectors and their hosts, thus interfering with the epidemiology of vector-borne diseases.

Evidence coming from different species of haematophagous insects, including mosquitoes and bedbugs, shows that the quick intake of a large quantity of fluid much warmer than the arthropod itself is associated to thermal stress (Benoit et al. 2011). Different mechanisms are activated at each feeding event, to counteract the deleterious effects associated with the ingestion of a warm blood meal and the risk of overheating: 1) the rapid synthesis of Heat Shock Proteins (HSPs) and 2) the activation of thermoregulatory mechanisms (Benoit et al. 2011, 2019; Lahondère and Lazzari, 2012; Paim et al. 2016; Lahondère et al. 2017). Even though insects and acarians certainly differ in their anatomical and physiological characteristics, we hypothesize that they are both exposed to thermal stress during blood feeding. The question arises then, about how they avoid the potential deleterious effects associated with feeding on warm-blooded vertebrates. It has been shown that several HSPs, including HSP70 and HSP90, are greatly upregulated in the *Ornithodoros moubata* (Murray, 1877) midgut after feeding (Oleaga et al., 2017). Moreover, in hard ticks, it is known that HSPs are overexpressed during feeding and that they play a crucial role in the mechanisms evolved by ticks in order to survive during blood feeding and to tolerate pathogen infection (Guilfoile and Packila, 2004; Oleaga et al., 2017).

To investigate whether soft ticks have developed any thermoregulative strategy to avoid overheating during blood-feeding, we analysed the evolution of the body temperature of ticks while feeding on a host, using real-time infrared thermography. Based on these results, we then evaluated the role that the cuticle wrinkled architecture could plays in thermoregulation in the soft tick *O. rostratus*.

## 2. Material and methods

### 2.1 Ticks

*Ornithodoros rostratus* (Aragão, 1911) ticks used in the experiments have been reared in the laboratory for five years from specimens collected in Nhecolândia, Mato Grosso do Sul, Brazil (19° 03’ S, 56° 47’ W) (Costa et al., 2015). Ticks were maintained inside an incubator at 28±2°C temperature and 85±10% humidity. They were fed on Swiss mice (*Mus musculus*) every 20 days.

### 2.2 Blood-feeding assays and thermography

Fourth instar nymphs, starving from 20 to 30 days, were fed on an anaesthetised hair-less mouse (150 mg kg^−1^ ketamine hydrochloride, Cristalia, Itapira, SP, Brazil and 10 mg kg^−1^ xylazine, Bayer S.A., São Paulo, SP, Brazil; intraperitoneally) placed over a heating pad, to avoid drug-induced hypothermia. The temperature of each tick and the host were simultaneously recorded during the whole feeding process by means of real-time infrared thermography. A thermographic camera (Pyroview, Dias Infrared Systems, Germany) equipped with a macro lens was used (Fig. 1, see Lahondère and Lazzari, 2012 for further details). This non-invasive procedure allowed recording thermographic videos, unravelling the dynamics of changes in the temperature of the tick across its body without disturbing it while feeding on the host. Videos were then analysed using the Pyrosoft software. All experiments were performed in a room maintained at 25°±1°C and 40 individual blood-feeding recordings were obtained.

**Figure 1.**
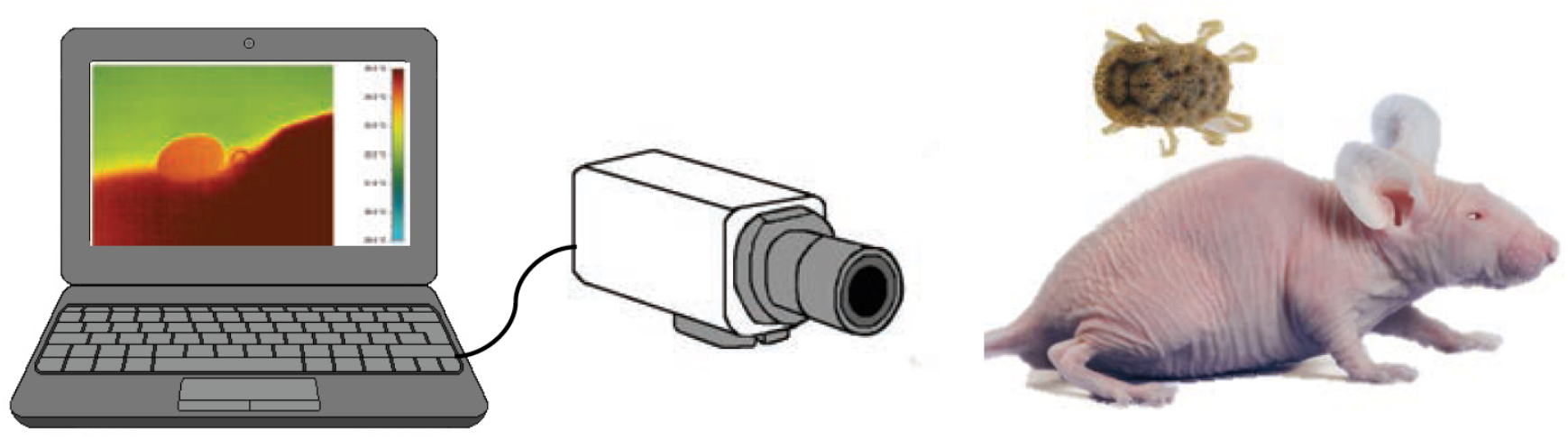
Experimental setup for real-time thermographic recording of the dynamics of temperature changes in an *Ornithodoros rostratus* nymph feeding on an anesthetised hairless mouse. A thermographic camera equipped with a macro lens, was connected to a laptop computer to record the evolution of the tick body surface temperature.

### 2.3 Fluorescein assays

In order to analyse to what extent the architecture of the tegument could influence heat dissipation, we performed an experiment in which we applied drops (approximately 10 uL) of a fluorescent solution composed of 1% fluorescein (Sigma-Aldrich) in saline solution on the cuticular surface of ticks (N = 5), and monitored its spread by observing its fluorescence under UV-light excitation using an epifluorescence microscope and low magnification.

All procedures were done in accordance with the manuals for experiments using animals and were approved by the Ethics Committee for the Use of Animals (CEUA, Instituto de Ciências Biológicas, Universidade Federal de Minas Gerais) under the protocol 301/2013.

## 3. Results

### 3.1 Feeding on a live host

Once placed on the anesthetised mouse, ticks inserted their mouthparts in the host skin and remained immobile. Not all individuals could be followed from an initial biting to full engorgement. Some ticks fed only partially and left the host; others detached and bit again and, in a few cases, movements did not allow us to keep a proper focus with the thermographic camera, which lead to a decrease in reliable temperature measurements. Engorgement was noted when the tick’s body became gradually rounder. After 8.2±1.7 min (mean±s.e.m., n= 7), a drop of fluid appeared on the ventral part of the tick and became gradually larger, due to the continuous secretion of the coxal glands (Lavoipierre and Riek, 1955; Kaufman et al. 1981, 1982). This fluid continued flowing until the end of the feeding.

### 3.2 Blood-feeding assays and thermography

When a tick was over the skin of a mouse, given its small body mass and the close contact surface (the whole ventral side), the temperature of the tick quickly increased to a value close to the temperature of its host (34°-35°C). When feeding started, the body temperature either slightly increased with blood intake or remained relatively constant (Fig. 2). When fluid excretion began, the temperature of the tick gradually decreased to reach a minimum by the end of feeding, around 3°C lower than the initial value. (Fig. 3, Supp. Video).

**Figure 2.**
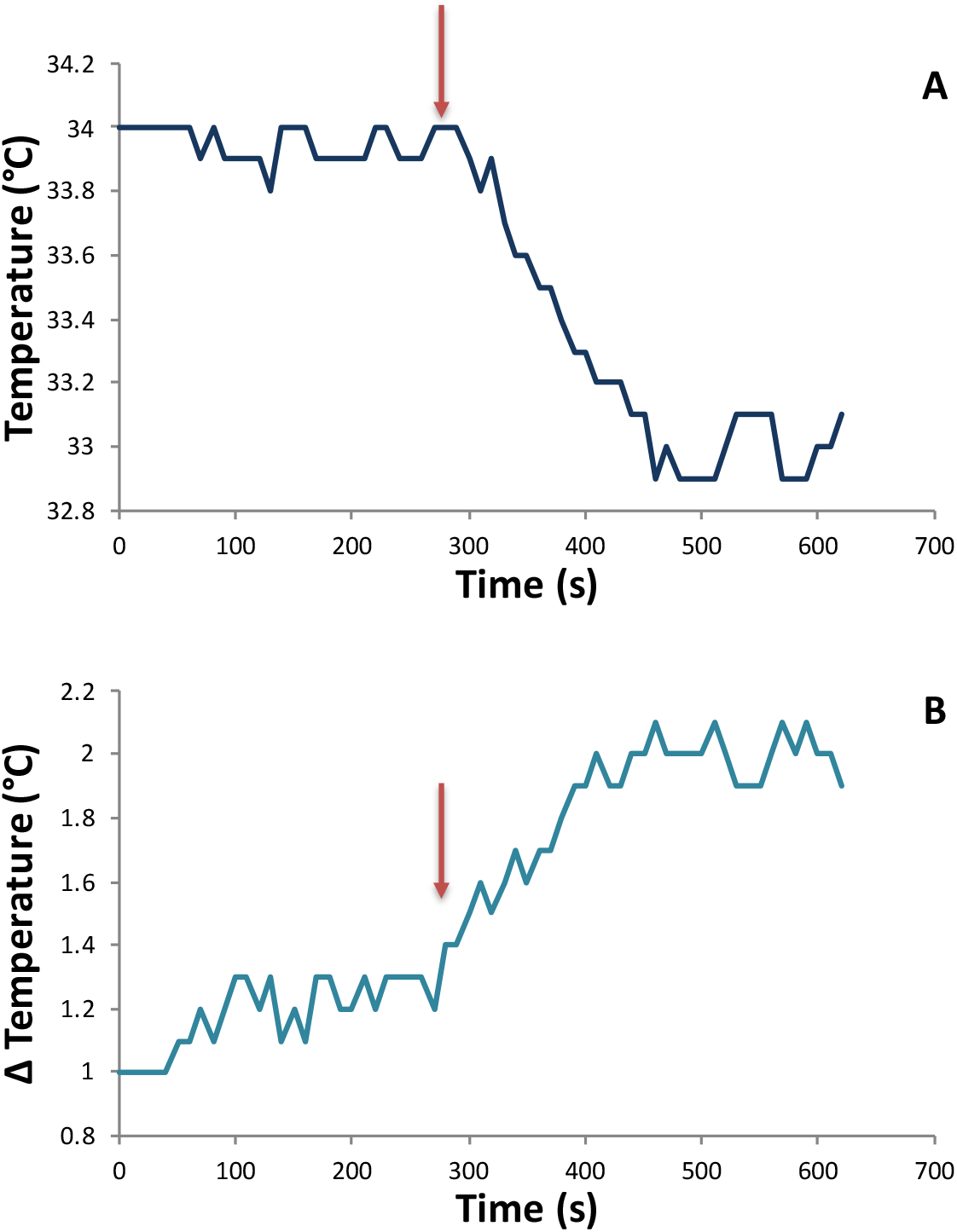
Sample record of the temperature of a tick during feeding. **A.** Temperature on the tick body surface. **B.** Temperature difference between the tick and the mouse. Red arrows indicate the moment when the secretion of coxal fluid began.

**Figure 3.**
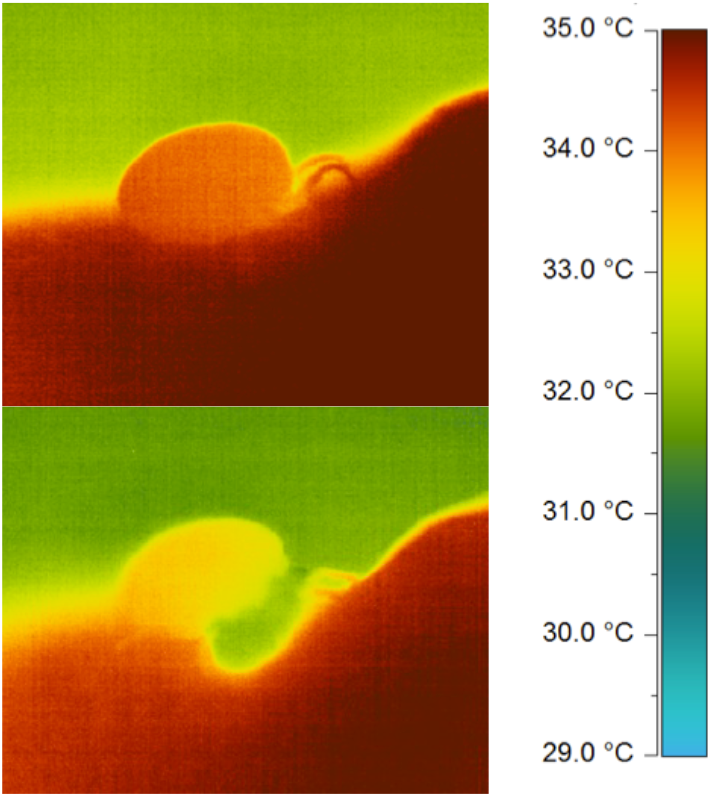
Thermographic images of a tick, before fluid secretion (top) and during fluid secretion (bottom). Different colours indicate different temperatures, as indicated in the reference scale on the right. A drop of coxal fluid (in green) can be observed in the ventral region of the tick in the bottom image.

The Figure 4 depicts the difference between the temperature of the tick and that of the host, before and after the secretion of fluid began. We found that the excretion of fluid led to a significant decrease of the tick body temperature (Fig. 4).

**Figure 4.**
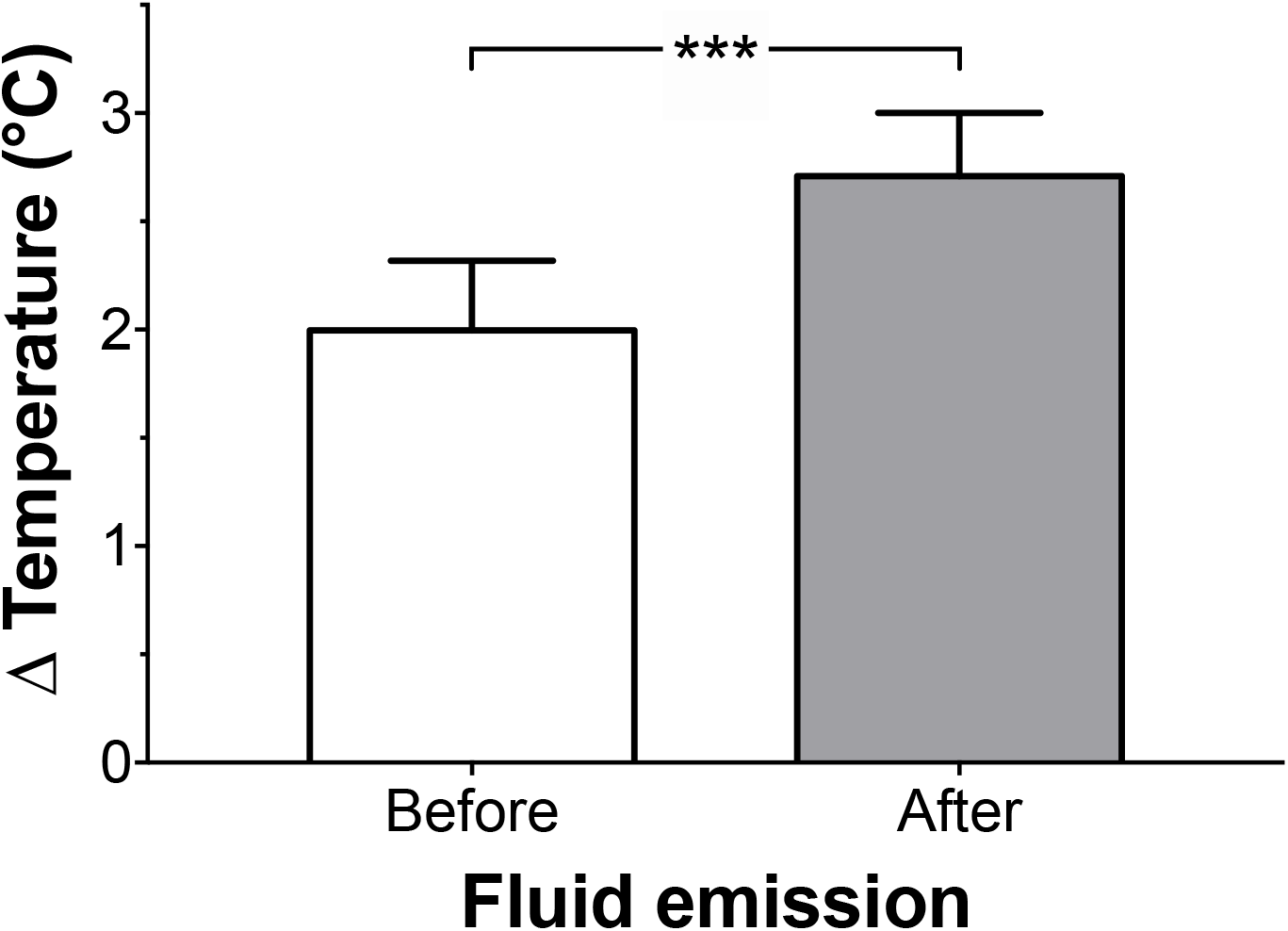
Temperature difference between the tick and the host before and after the secretion of coxal fluid began. The statistical comparison rendered a highly significant difference (paired Student *t*-test, p<0.01, n= 7).

### 3.3 Fluorescein assays

Given the particular architecture of the tegument of *Ornithodoros* along with the synchrony between secretion of fluid and temperature decrease, we suspected that a process of evaporative cooling could be involved. We also hypothesised that it would be facilitated by capillary diffusion through the three-dimensionally anfractuous structure of *Ornithodoros* cuticle (*e.g.* Labruna et al., 2016; Muñoz-Leal et al. 2016, 2017). Besides, the fact that temperature changes were observed to be simultaneous and similar over the whole body (*i.e*., no heterothermy was observed, Fig. 3), suggested that fluid evaporation could occur over the whole tegument surface.

To test this hypothesis, we observed the dynamics of spread of a fluorescent solution in different individuals (n = 5). Figure 5 highlights two images of the same individual taken 30s apart, illustrating the rapid increase of the fluorescent area by capillarity, and representative of what was observed in all tested individuals.

**Figure 5.**
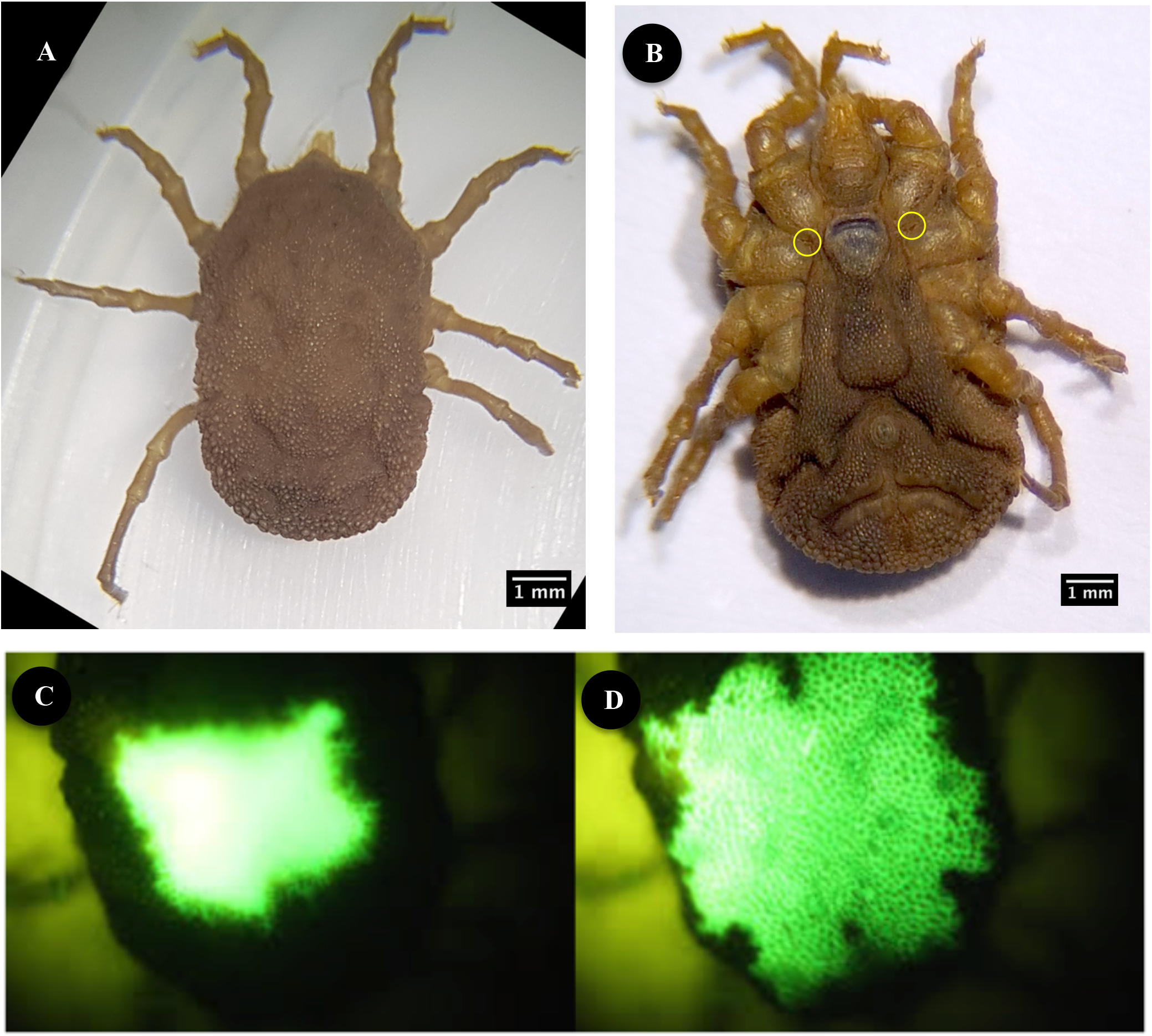
**A**. Dorsal and **B**. Ventral views of an adult *Ornithodoros rostratus*. Circles indicate the opening of coxal glands. C-D. Sample record of the spread of a drop of fluorescein solution over the dorsal cuticular surface of a tick, immediately after dropping (left) and 30s later (right).

We can then conclude that the spread of the excreted fluid over the cuticular surface lead to an increase of the evaporative surface, thus facilitating the cooling of the acarian during blood-feeding.

## 4. Discussion

The fact that haematophagous arthropods are exposed to thermal stress at each feeding event was only recently revealed, along with the existence of molecular recovery strategies and thermoregulatory mechanisms to avoid overheating during blood-feeding (Benoit et al. 2011, 2019; Lahondère and Lazzari, 2012; Lahondère et al. 2017).

Among acarians, the synthesis of HSPs induced by heat shocks and blood-feeding in has been characterised in the hard tick *Ixodes scapularius* (Guilfoile and Packila, 2004; Busby et al. 2012), and more recently in the soft tick *O. moubata* (Oleaga et al. 2017). A functional relationship between thermal stress during feeding and HSP synthesis can thus be hypothesized to be a general mechanism in blood-sucking arthropods associated to endothermic hosts (Pereira et al. 2018).

Until now, thermoregulatory mechanisms associated with blood-feeding were only known to exist in insects (Lahondère and Lazzari, 2012, Lahondère et al. 2017). Our present study in *O. rostratus* not only highlighted a new ectothermic species possessing thermoregulatory abilities, suggesting that it may be also present in other acarians, but also revealed a new original mechanism, based on morphological and physiological specific adaptations. The location and intense activity of the coxal gland during feeding, as well the cuticular anfractuosities of the tegument of *Ornithodoros* ticks, both contribute to the rapid dissipation of heat excess. The spread of the fluid over a large tegument surface, facilitates the heat exchange between the fluid and the body, as well as the evaporation of the fluid, reducing the body temperature of the tick by evaporative cooling. Interestingly, Yoder et al. (2009) have shown that dermal gland (*sensilla sagittiformia*) secretions in the brown dog tick *Rhipicephalus sanguineus* are implicated in heat tolerance. Indeed, the secretion and spread of this liquid on the cuticle allow this species to avoid heat stress while feeding on their host. As the exact way this gland secretion is involved in heat tolerance remains unclear, testing for the hypothesis of evaporative cooling during blood-feeding in this species could shed some light on the fine mechanisms underlying the role of these secretions in heat tolerance in the *R. sanguineus* ticks. It has also been suggested that the folded architecture of the *Ornithodoros* ticks is a derived character from their ancestor, *Nuttalliella namaqua* allowing the tick to greatly expand its abdomen and ingest an important quantity of blood quickly (Klompen and Oliver, 1993; Mans *et al.* 2011). This species is lacking coxal glands but instead possesses an anal pore, through which excretion occurs. Whether these ticks might use evaporative cooling during feeding remains unknown, but would be interesting to investigate in order to have a better understanding of the evolution of thermoregulatory processes among blood-sucking insects and acarians.

This is the first time that a thermoregulatory mechanism is unravelled in acarians. Among insects, evaporative cooling has been shown to occur in several species including aphids (Mittler, 1958), honeybees (Heinrich, 1979; 1980) and moths (Adams and Heath, 1964). Honeybees and moths regurgitate droplets of nectar that they maintain on their mouthparts to cool down their head thus avoiding overheating due to either high ambient temperature or activation of their flight thoracic muscles (*i.e.* endothermy) (Adams and Heath, 1964; Heinrich, 1979). Moreover, honeybees move the droplets in and out of their mouth to cool down their head more efficiently (Heinrich, 1980). They are thus able to decrease their head temperature by 3°C and 4°C below ambient temperature, respectively. Aphids cool down their abdomen by excreting honeydew and by keeping a typical posture (*i.e.* away from the substrate) which increases air movement around their abdomen (Mittler, 1958). Among blood-sucking insects, *Anopheles* mosquitoes are known to use droplets composed of urine and undigested blood to cool down their abdomen during feeding (Lahondère and Lazzari, 2012). Soft ticks ingest blood up to 12-fold their own body weight and to restore their water balance as well as to concentrate the nutrients in the ingested blood, they excrete large amount of coxal fluid which is hypoosmotic. This fluid, unlike anopheline mosquitoes which might also excrete fresh erythrocytes (Briegel and Rezzonico, 1985), is mainly composed of water, electrolytes, nucleic and amino acids, proteins, carbohydrates and lipids (Frayha et al., 1974). It is worth mentioning that coxal fluid is excreted by soft ticks when exposed to temperatures over 40°C (Kaufflan and Sauer, 1982; Rémy, 1922) thus reinforcing the hypothesis that coxal fluid excretion participates in thermal stress management in these ticks.

Finally, evaporative cooling is not the sole mechanism for thermoregulation that has been described in haematophagous insects. Indeed, the kissing bug, *Rhodnius prolixus*, possesses in its head an elaborated counter-current heat-exchanger, which reduces the temperature of the ingested blood, before it reaches the abdomen of the bug (Lahondère et al., 2017). It is then possible, that other tick species could have developed other strategies for reducing thermal stress during blood feeding.

## 5. Conclusions

Our study shows for the first time in acarians, that soft bodied ticks perform thermoregulation during feeding by excreting fluids through their coxal gland that spreads to their cuticle and cool them down. These results extend previous knowledge obtained in insects, to other blood-sucking arthropods. We have just began disentangling how blood-sucking arthropods manage avoiding the deleterious effects associated with feeding on endothermic vertebrates, and much work remains to be conducted to get a full picture of the spectrum of possible adaptations developed by these arthropods during their long evolutionary history.

## Author contribution

Experimental design: C.R.L, C.L. and M.H.P.

Thermographic recordings: C.R.L.

Data Analysis: A.F. and C.R.L.

Tick rearing and expertise: R.N.A.

Fluorescent microscopy: M.H.P.

Manuscript draft: C.R.L.

Final elaboration and editing of manuscript: all the authors

## Acknowledgements

We are deeply grateful to the technical staff of the Brazilian laboratory for their valuable assistance with experiments. This work of C.R.L. in Brazil was possible thanks to the financial generous support of the Brazilian program “Science without borders” (CNPq). The work was supported by the CNRS and the University of Tours (France) and by the Conselho Nacional de Desenvolvimento Cientı́fico e Tecnológico (CNPq) and Instituto Nacional de Ciência e Tecnologia em Entomologia Molecular (Brazil). The work of A.F. was conducted in the frame of her second-year master studies at the University of Tours (France).

## Supplementary material

**Movie 1.** Example of a thermogram of an *Ornithodoros rostratus* nymph during blood-feeding on a hairless mouse. At the beginning of the process, the temperature of the thick increases as a consequence of blood ingestion. Then, the accumulation of fluid can be observed in the anterior (right) part of the body, followed by a decrease in the temperature of the acari. The sequence has been accelerated, in order to present a process usually taking around 30 min.

